# Ribozyme-Mediated, Multiplex CRISPR Gene Editing and CRISPRi in *Plasmodium yoelii*

**DOI:** 10.1101/481416

**Authors:** Michael P. Walker, Scott E. Lindner

## Abstract

Functional characterization of genes in *Plasmodium* parasites often relies on genetic manipulations to disrupt or modify a gene-of-interest. However, these approaches are limited by the time required to generate transgenic parasites for *P. falciparum* and the availability of a single drug selectable marker for *P. yoelii*. In both cases, there remains a risk of disrupting native gene regulatory elements with the introduction of exogenous sequences. To address these limitations, we have developed CRISPR-RGR, a SpCas9-based gene editing system for *Plasmodium* that utilizes a Ribozyme-Guide-Ribozyme (RGR) sgRNA expression strategy. Using this system with *P. yoelii*, we demonstrate that both gene disruptions and coding sequence insertions are efficiently generated, producing marker-free and scar-free parasites with homology arms as short as 80-100bp. Moreover, we find that the common practice of using one sgRNA can produce both unintended plasmid integration and the desired locus replacement editing events, while the use of two sgRNAs results in only locus replacement editing. Lastly, we show that CRISPR-RGR can be used for CRISPR interference (CRISPRi) by binding dCas9 to targets in the gene control region of a gene-of-interest, resulting in a modest reduction in gene expression. This robust and flexible system should open the door for in-depth and efficient genetic characterizations in both rodent- and human-infectious *Plasmodium* species.

**Importance:** *Plasmodium* parasites, the causative agent of malaria, still pose an enormous threat to public health worldwide. Gaining additional insight into the biology of the parasite is essential for generating an effective vaccine and identifying novel drug targets. To this end, CRISPR/Cas9 tools have been developed to more efficiently interrogate the *Plasmodium* genome than is possible with conventional reverse genetics approaches. Here, we describe CRISPR-RGR as an addition to the CRISPR/Cas9 toolbox for the rodent-infectious *Plasmodium* parasites. By using multiple ribozyme-flanked single guide RNAs expressed from RNA polymerase II promoters, transgenic parasites can be rapidly generated as designed without leaving selectable markers. Moreover, CRISPR-RGR can be adapted for use as a CRISPR interference (CRISPRi) system to alter gene expression without genome modification. Together, CRISPR-RGR for gene editing and CRISPRi application can hasten investigations into the biology and vulnerabilities of the malaria parasite.

## Introduction

Malaria remains one of the world’s most daunting public health concerns, with over 200 million infections and nearly half a million fatalities every year (WHO, 2017). Despite gains made to reduce transmission worldwide, there is still a need for a highly effective, licensed vaccine and additional anti-malarial drugs to respond to and overcome the emergence and spread of drug resistance. To produce these new therapeutics, further studies of the causal agent of malaria, the *Plasmodium* parasite, are required. These studies typically rely upon reverse genetic techniques to disrupt or tag genes-of-interest, however, producing gene modifications in *Plasmodium* parasites has inherent challenges. First, transfection efficiencies are surprisingly low compared to that of model eukaryotes and human cells, despite advances in nucleofection technologies improving these efficiencies a thousand fold over initial methods (1). Selection of transgenic parasites commonly takes months to achieve locus replacement events in *P. falciparum* through the use of both positive and negative drug pressure. Moreover, only a single drug selectable marker (DHFR) is available for modifications of the genome of rodent-infectious species of *Plasmodium*, such as *P. yoelii*, *P. berghei*, and *P. chabaudi* (2-4). In response to this, several methodologies have been developed to recycle DHFR expression cassettes (Gene In Marker Out (GIMO), FLP/Frt and CRE/lox recombination-based methods), but these methods are time consuming and require multiple interventions (5-7). Finally, exogenous sequences are essentially always left in the parasite genome, which can include expression cassettes that can influence transcription of neighboring genes, plasmid backbone sequences, and in the best cases, a single frt or lox site following recombinase excision (8-10).

The recent development of the Clustered, Randomly Interspaced Short Palindromic Repeats/CRISPR-associated protein 9 (CRISPR/Cas9) gene editing system has been shown to improve the efficiency of genome editing. This has been observed for both *Plasmodium falciparum*, the human-infectious species responsible for the majority of cases and deaths, and *Plasmodium yoelii,* a rodent-infectious species typically favored for rapid genetic manipulation, the ability to interrogate the entire life cycle, and similarities with *P. falciparum* in mosquito stage development and host-pathogen interactions (11-15). Importantly, CRISPR-based gene editing doesn’t rely on integration of a selection cassette into the genome, and the edited genome can therefore be free of drug resistance/fluorescent markers and scar sequences. CRISPR-based gene editing therefore allows the interrogation of multiple genes either simultaneously or sequentially, and if properly controlled, could be done at multiple points throughout the parasite’s life cycle.

CRISPR/SpCas9 gene editing systems function by expressing sgRNAs (single guide RNAs) that recruit the *Streptococcus pyogenes* Cas9 (SpCas9) endonuclease to a complementary 20nt sequence of genomic DNA. There, a double-stranded break (DSB) is created and typically repaired with either the error-prone Non-Homologous End Joining (NHEJ) or by Homology-Directed Repair (HDR) pathways. Because *Plasmodium* parasites lack several essential proteins required for NHEJ, homology-directed repair of double-strand breaks predominates, with infrequent repair also occurring by microhomology-mediated end joining (MMEJ) (16, 17).

Therefore, strategies for CRISPR gene editing in *Plasmodium* require the introduction of three components: SpCas9, sgRNAs, and a homology directed repair (HDR) template, which are typically encoded on one or two nuclear plasmids.

CRISPR-based gene editing of *P. falciparum* has been achieved using dual-plasmid systems, with each plasmid encoding a unique drug resistance marker, and either SpCas9 or the HDR and sgRNAs (11, 12). Efficient expression of sgRNAs was demonstrated with either an engineered T7 RNA polymerase system or with the RNA polymerase III transcribed *P. falciparum* U6 promoter (11, 12). Upon electroporation, parasites are then pressured with one or both drugs simultaneously to select for parasites containing all of the necessary gene editing elements (11, 12).

CRISPR/Cas9 strategies in *P. yoelii*, however, are limited by the availability of only one drug-selectable marker (DHFR), so all CRISPR/SpCas9 gene editing elements must be packaged onto a single vector to allow their selection. Previously described single-plasmid systems use the *P. yoelii* U6 promoter to express sgRNAs, however the resulting transgenic parasites were found to retain the plasmid sequences and remained resistant to drug pressure (13). Because of this, a second system was constructed that included a negative-selectable marker in the plasmid backbone, so that parasites that retained a nuclear plasmid or that may have integrated the plasmid could be selected against following gene editing (18). Despite these limitations, the laboratory of Jing Yuan has used CRISPR to methodically interrogate the ApiAP2 gene family and genes important for ookinete motility, and have created transgenic parasites that express a constitutively expressed SpCas9 nuclease, or male and female-enriched fluorescent protein reporters (13, 18-21).

Although significant progress has been made to date in the development of CRISPR tools for use in *Plasmodium*, significant limitations remain. First, existing systems use RNAP III promoters for sgRNA expression, which is preferred due to the well-defined 5’ and 3’ ends on this class of transcript. However, as RNAP III promoters are strong and constitutively active as required to produce 5S rRNA, tRNAs and other critical non-coding RNAs, their use in transcribing sgRNAs would not permit stage-specific or readily tunable expression. Secondly, most studies to date have targeted a single locus with one sgRNA, and those few efforts that have used multiple sgRNAs used multiple RNAP III-based cassettes to express the sgRNAs. Lastly, the adoption and use of nuclease-dead variants of SpCas9 (dSpCas9), which has been used in prokaryotes and eukaryotes for gene regulation by CRISPR activation (CRISPRa) and CRISPR interference (CRISPRi), has not been demonstrated. Because *Plasmodium* parasites lack genes essential for RNA interference (RNAi) (22), genetic tools to regulate transcript abundance (e.g. promoter swap, glmS) or protein abundance (e.g. DD/Shield1, EcDHFR-DD/Trimethoprim, TetR/DOZI) have been developed and utilized (23-26). However, these all require single or multiple modifications to the genome and introduce exogenous sequences into the locus-of-interest. CRISPRi is therefore a desirable tool, as it would allow for regulation of the expression of specific genes without modification of the genome itself.

Here, we show that CRISPR-RGR is able to effectively generate gene deletions, tag insertions, and can be used for CRISPR interference. Using a Ribozyme-Guide-Ribozyme (RGR) method of sgRNA expression, we show that three RNAP II promoters can be used to express multiple sgRNAs simultaneously, and that genome editing can be rapidly and efficiently achieved in *P. yoelii*. Additionally, we demonstrate that the number of sgRNAs used to target a gene influences the outcome of genome repair, and that negative selection is not required to produce parasites with only locus replacement events observed. Finally, we demonstrate that CRISPRi is possible in *P. yoelii*, with the efficacy of 11 individual sgRNAs targeting across the upstream portion of the gene control region of PyALBA4 demonstrating positional, but not strand-specific, effects. The contribution of CRISPR-RGR to the growing *Plasmodium* CRISPR toolbox explains and solves previous issues with current strategies that use one sgRNA, improves how synthetic biology approaches can be used to expedite gene editing, and provides a framework for CRISPRi-based gene regulation.

## Results

### Single-plasmid, ribozyme-mediated CRISPR/SpCas9 Plasmid Design for Rodent-Infectious *Plasmodium* Parasites

To streamline CRISPR/SpCas9 editing in *Plasmodium yoelii* parasites, we developed a flexible, single-plasmid construct that contains all necessary CRISPR/SpCas9 gene editing elements (Fig. 1, 1^st^ Generation). Expression of SpCas9::GFP, HsDHFR (to provide resistance to anti-folate drugs), and sgRNAs were generated by individual iterations of the strong, constitutive *pbef1α* promoter and the *pbdhfr* 3’UTR/terminator. Each of these cassettes is flanked by unique restriction enzyme sites for easy modification and substitutions. In addition to these cassettes, we incorporated a homology-directed repair (HDR) template to enable homology directed repair of the double-strand breaks (DSBs) that are created by SpCas9. Because the sequence conservation of these gene control elements between *P. yoelii* and *P. berghei* is exceptionally high, and that in fact a mixture of elements from both species are used in the plasmids describe here, we anticipate that gene editing using these plasmids will be possible in *P. berghei* as well.

**Figure 1:**
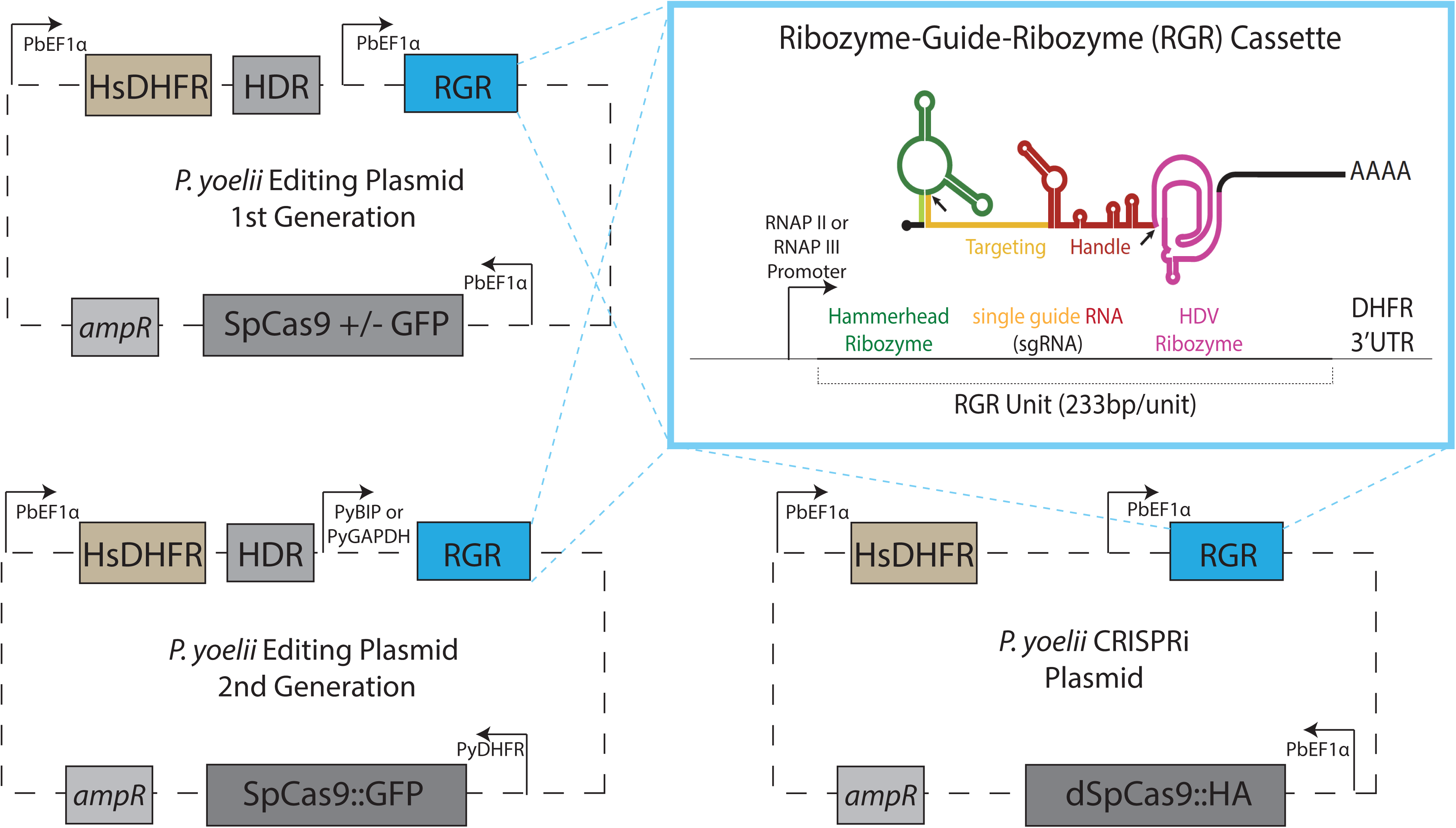
Single-Plasmid, Ribozyme-Based CRISPR/SpCas9 constructs for *Plasmodium yoelii* parasites. Each plasmid contains expression cassettes for SpCas9, a HsDHFR drug resistance marker, and sgRNA expression (see inset). For gene editing plasmids, a homology-directed repair (HDR) template is also included. Inset: Precise autocatalytic cleavage by both Hammerhead and HDV ribozymes release the sgRNA without extra bases attached. The location of the ribozyme cleavage is noted by the arrows.

The major differences between this CRISPR-based editing system for *Plasmodium* and those previously described lie in the expression of the sgRNAs and the preparation of the sgRNA and HDR template sequences. Existing *Plasmodium* CRISPR systems use RNA Polymerase III-driven U6 promoters or T7 RNA Polymerase-based systems for sgRNA expression (11, 12). In contrast, we have expressed a transcript encoding a Ribozyme-Guide-Ribozyme (RGR) unit that uses a minimal Hammerhead Ribozyme and a Hepatitis Delta Virus ribozyme to flank the sgRNA on its 5’ and 3’ ends, respectively (Fig. 1, inset). This RGR approach, first described in yeast (27) and since used in *Leishmania* (28) and zebrafish (29), generates sgRNAs with precisely defined 5’ and 3’ ends, and allows for the simultaneous expression of multiple guides under control of a single promoter, including RNAP II promoters. While RNAP II promoters are far more abundant than RNAP III promoters, they are not typically used for sgRNA expression as their transcripts can initiate from multiple transcriptional start sites (TSS’s) and are capped and polyadenylated. The potential impact of these 5’ extensions and modifications upon sgRNA activity are not sufficiently understood to confidently use them for this application. However, the inclusion of autocatalytic, self-cleaving ribozymes within an RNA eliminates these potential problems with RNAP II transcripts, and their beneficial properties of stage-specific and tunable expression levels can be used. Moreover, these individual RGR units can be polymerized into an RGR array on a single transcript and be used to generate multiple sgRNAs. In addition, our sgRNA design contains an extended duplex and a single base change compared to the original sgRNA sequence, which has been shown to have increase editing efficiency (30). Finally, we leverage advances in synthetic biology to create custom DNA fragments that can include the RGR transcript, short HDR templates, or both, which greatly expedites plasmid generation. We anticipate that as the cost of gene synthesis continues to decrease, these approaches can be scaled for use in both forward and reverse genetic screens. A step-by-step tutorial for construct generation is provided in Supplemental File 1.

### Genetic Disruption of *pyalba4*

In order to functionally test this single-plasmid, CRISPR-RGR system, we targeted the gene encoding the PyALBA4 RNA-binding protein, which we have previously characterized (10). Using conventional reverse genetics approaches, we have shown that *pyalba4* can be deleted in asexual blood stage parasites, which results in the production of 2-3 fold more mature male gametocytes that can exflagellate as compared to wild-type parasites. Additionally, a C-terminal GFP tag can be introduced with no observable effect upon parasite growth or transmission. In order to delete *pyalba4* by CRISPR-RGR, two sgRNA targets were chosen at the 5’ and 3’ ends of its coding sequence by manually scanning for NGG PAM motifs and subsequent computational assessment using the Eukaryotic Pathogen CRISPR guide RNA/DNA design tool (http://grna.ctegd.uga.edu) (Fig. 2A, red vertical lines). This tool provides a score for each sgRNA based on the target specificity within the genome, as well the GC content and position-specific nucleotide composition for bases that have been shown to affect sgRNA efficiency. Additionally, this tool will flag any sgRNA with long poly-T tracts (more than four in a row) which can cause early termination of RNAP III transcripts. Because the RGR system utilizes RNAP II promoters, we are not limited by sgRNAs containing poly-T tracts that are prevalent in *Plasmodium* genomes (31). The initial HDR template was designed with homology arms comprised of ∼800bp of sequence homologous to the target gene on either side of the two DSBs, with a unique 18bp DNA barcode between them that could be used for unambiguous, simple genotyping PCR. Notably, this barcode is not necessary for genome editing and could be omitted to produce completely scar-less modifications without the introduction of any exogenous sequences.

**Figure 2:**
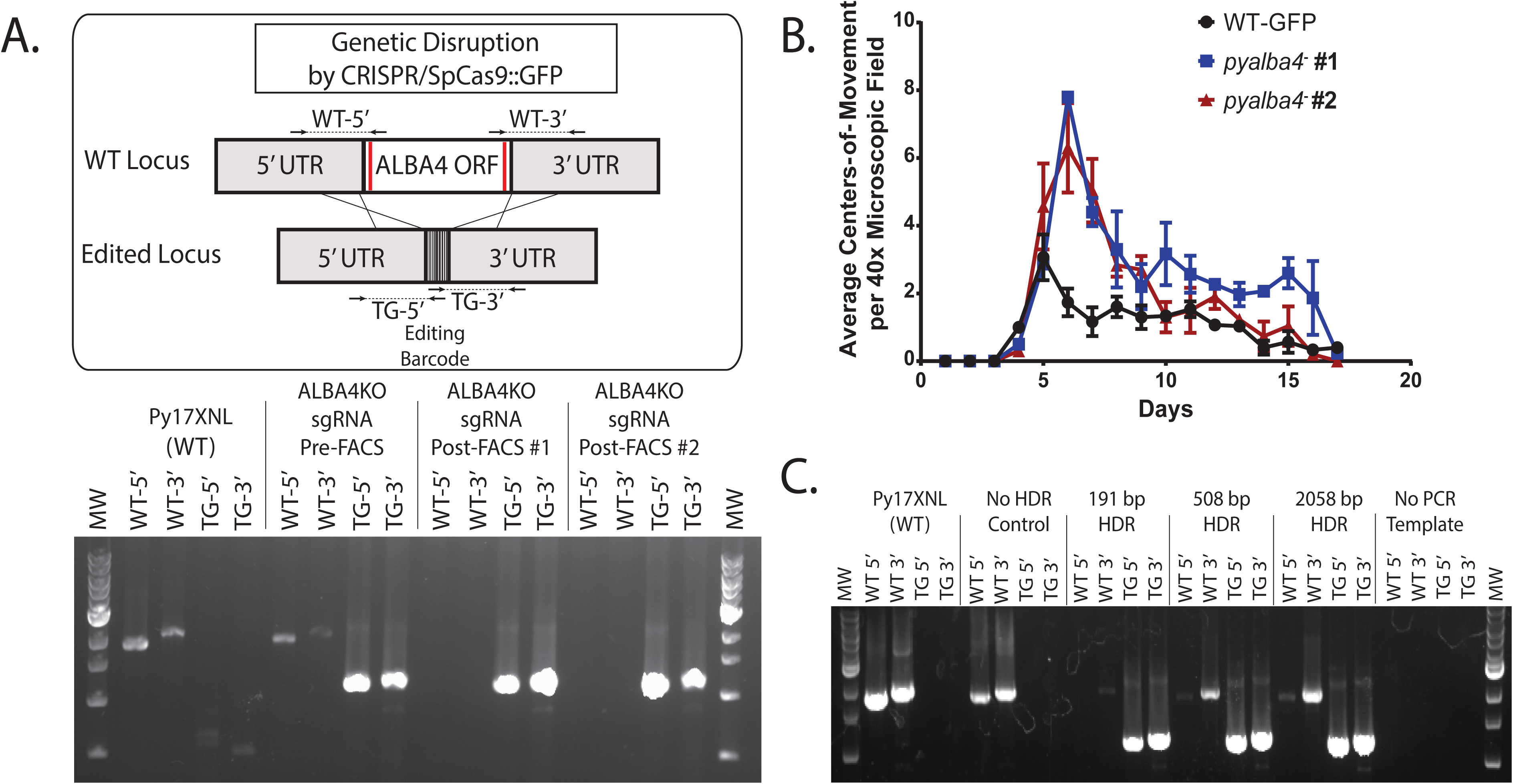
A Single Plasmid, Ribozyme-based CRISPR/SpCas9 system efficiently edits the *P. yoelii* genome. A.) The *pyalba4* ORF was deleted by expressing two sgRNAs that target the 5’ and 3’ ends of the coding sequence, with transgenic parasites seen within six days of constant drug selection. Genotyping PCR showed that the majority of the parasite population had been edited by locus replacement. Enrichment by FACS of parasites expressing SpCas9::GFP resulted in the selection of a completely transgenic parasite population by genotyping PCR. B) Two clones of the CRISPR-generated *pyalba4*^−^ transgenic parasites produce 2-3 fold more male gametocytes that can activate into gametes as compared to wild type parasites. C.) A range of HDR template sizes were tested (191, 508, 2058bp). All HDR templates allowed efficient editing of the *pyalba4* locus, with the smallest (about 80-100bp of homologous sequences on either side of the DSB’s) showed the most efficient repair by genotyping PCR.

Upon transfection of this plasmid into wild type (WT) *Plasmodium yoelii* (17XNL strain) parasites with constant pyrimethamine selection, mice reached 1% parasitemia in only 8 days (2). We observed that a large subset of these parasites showed expression of SpCas9::GFP by live fluorescence microscopy (data not shown), and genotyping PCR analysis showed efficient editing of the *pyalba4* locus (Fig. 2A bottom). Furthermore, by enriching for SpCas9::GFP-positive schizonts via Fluorescence-activated Cell Sorting (FACS), only edited parasites were present by genotyping PCR. Thus, this CRISPR-RGR approach rapidly produced a transgenic parasite population with no observable wild-type parasites present using as few as two mice.

We further verified that this population of CRISPR-generated *pyalba4*^−^ parasites had the same phenotype as *pyalba4*^−^ transgenic parasites generated by conventional reverse genetic approaches. Quantification of the activation of male gametocytes into gametes (measured as centers-of-movement/exflagellation centers via light microscopy) in the *pyalba4*^−^ line revealed a similar 2-3 fold increase in the number of activated male gametocytes as compared to wild type parasites, which was sustained across the full duration of the infection (Fig. 2B).

Because CRISPR-RGR rapidly and efficiently produced transgenic parasites when providing large homology arms in the HDR template, we tested the effect that lengthening (∼1000bp each arm) and shortening (∼80-100bp, ∼250bp each arm) the homology arms had upon gene editing. We observed that all homology arm lengths allowed for efficient gene editing, and that the smallest HDR tested (80-100bp each arm) had the most efficient editing (as evidenced by the least intense PCR amplicons for wild-type parasites) and could be selected in the same amount of time as was required for the longer HDR templates (Fig. 2C). Importantly, HDR arms of this length, even with the skewed A-T content *of P. yoelii*’s genome, can be chemically synthesized.

It is notable that over the course of these experiments, we observed that recombination was occurring in *E. coli* between two instances of the *pbef1α* promoter, and that the RGR portion of the plasmid was being excised. To stabilize the plasmid, we constructed a second generation of editing plasmids with no repeated elements and have not observed spurious recombination events occurring with this new design (Fig. 1 2^nd^ Generation, Supp. Fig. 1). This second generation design includes a single iteration of the *pbef1α* promoter and *pyef1α* 3’UTR to control expression of HsDHFR, the *pydhfr* promoter and *pbdhfr* 3’UTR to control expression of SpCas9::GFP, and either the *pybip* promoter or *pygapdh* promoter with the *pybip* 3’UTR to control transcription of the RGR element. Using the same sgRNA targets and the 191bp HDR template, we found that both promoters driving RGR expression edited parasites efficiently (Supp. Fig. 1). Furthermore, we again verified with FACS and genotyping PCR that parasites expressing SpCas9::GFP only contained the edited *pyalba4* locus. Together these results show that single-plasmid CRISPR-RGR is able to efficiently and robustly create gene deletions in *P. yoelii* parasites.

### Insertion of a GFP Reporter by CRISPR-RGR

We next used CRISPR-RGR to append a C-terminal GFP epitope tag to PyALBA4. To accomplish this, we chose a single sgRNA target (1x sgRNA) 18bp downstream of the stop codon and designed a HDR template with ∼700bp homology arms flanking an in-frame GFP tag (Fig. 3a). A base change of the NGG PAM sequence (shield mutation) was included in the HDR template so that SpCas9 ceases to bind and cut this locus post-editing. Transfection of this plasmid, along with constant pyrimethamine selection for 7 days, yielded a largely transgenic parasite population (Fig. 3B, left) with PyALBA4::GFP expression matching what was seen in transgenic parasites created by conventional methods (Fig. 3C).

**Figure 3:**
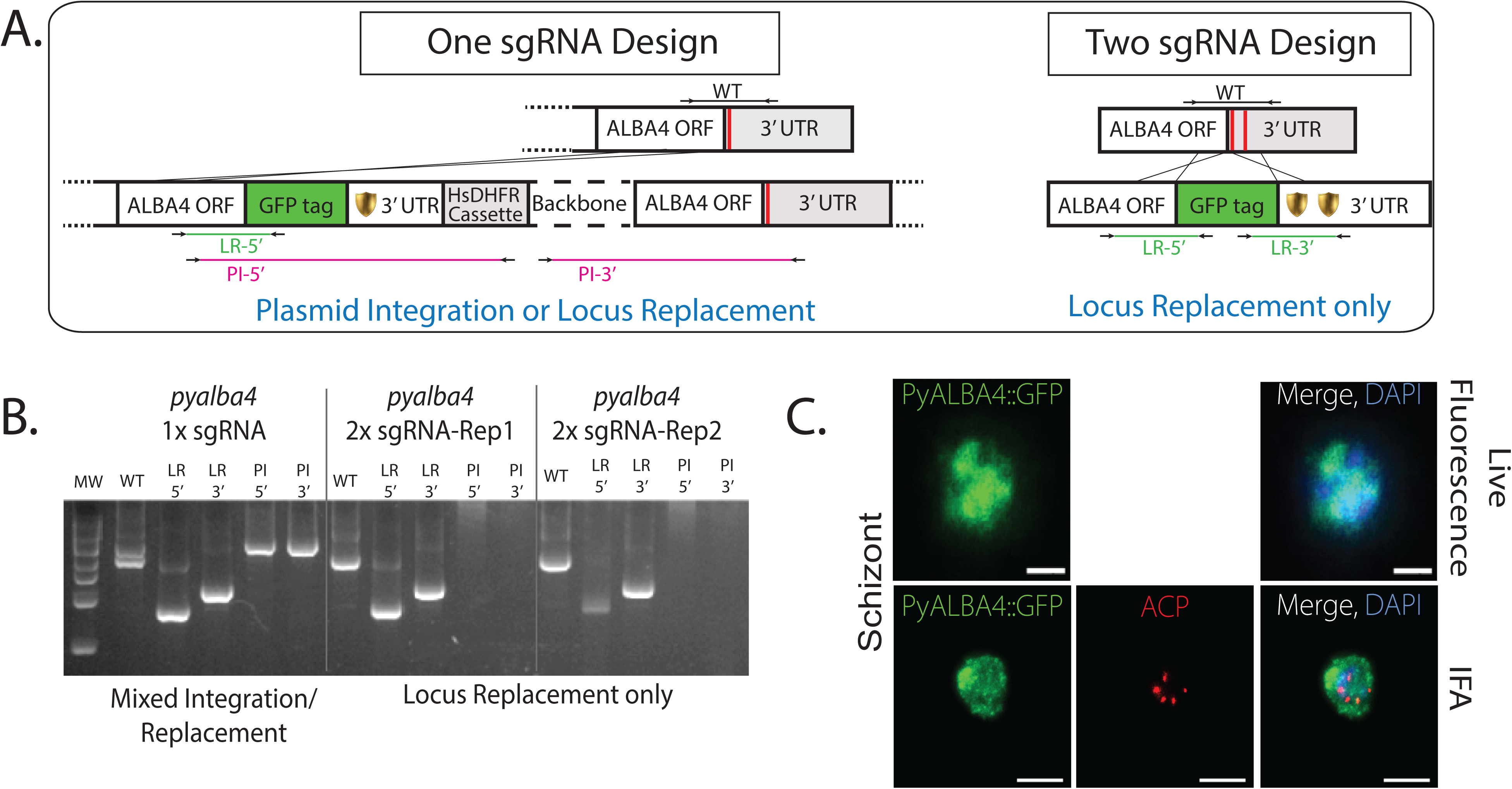
The number of sgRNAs used influences the outcome of genome repair. A.) Insertion of a GFP tag at the 3’ end of the *pyalba4* coding sequence was accomplished by using either one or two sgRNA targets (red lines in schematic). Each of these require a shield mutation in the HDR template to destroy the PAM site and prevent SpCas9 cleavage after the repair. B.) Genotyping PCR shows that the use of one sgRNA resulted in both plasmid integration into the targeted locus, as well as the desired locus replacement event. The use of two sgRNAs resulted in only locus replacement and no plasmid integration. C.) CRISPR-modified PyALBA4::GFP parasites show an expression pattern of ALBA4 in schizonts similar to that seen with conventionally-tagged PyALBA4 parasites. Scale bar = 5μm.

In contrast to conventional reverse genetic techniques, CRISPR/SpCas9 gene editing can be accomplished without leaving a drug selectable marker in the edited locus. This enables multiple, sequential gene edits to be made after curing the delivery plasmid, which is particularly useful with rodent-infectious malaria parasites where only one drug selectable marker (DHFR) is available. However, previous work has shown that completely curing these plasmids is challenging, as resistant parasites can be recovered more than 50 days post-removal of drug pressure (18). To circumvent this issue, methods have been developed to negatively select against parasites that retained the delivery plasmid following genome modification (18). We similarly attempted to cure the plasmid from FACS-selected PyALBA4::GFP parasites produced using one sgRNA, and observed that the parasites remained drug resistant after more than 14 days following removal of drug pressure. An expanded genotyping PCR assay that interrogates for both plasmid integration and locus replacement showed that both editing outcomes occurred (Fig. 3B).

We reasoned that introduction of two DSBs, and thus two genome repair events, would ensure that only locus replacement events would result. To test this, we selected a second sgRNA target in the *pyalba4* 3’ UTR downstream of the original sgRNA target, and introduced a second shield mutation into the HDR template (Fig. 3A). Transfection of this plasmid, coupled with constant selection with pyrimethamine, produced PyALBA4::GFP-expressing parasites. As before, genotyping PCR showed that a significant fraction of the parasite population had been modified. Notably, this two sgRNA design yielded only the desired locus replacement events and showed no evidence of plasmid integration (Fig. 3B, right). Furthermore, upon removal of drug pressure, the parasites quickly (<1 week) became sensitive to pyrimethamine once more and were amenable to subsequent transfections.

### A Single-Plasmid System for Ribozyme-Mediated CRISPR Interference

CRISPR activation (CRISPRa) and CRISPR interference (CRISPRi), which target a nuclease-dead variant of *S. pyogenes* Cas9 (dSpCas9) to locations on the genome to activate or repress transcription respectively, have emerged as effective methods to study essential genes in prokaryotes, model eukaryotes and human cells (32-34). CRISPRa and CRISPRi largely rely upon the fusion of transcriptional activation or repression domains to dSpCas9 to produce these effects on transcription. However, domains that typically function in most eukaryotes (e.g. VP16, KRAB) are not known to work in *Plasmodium*, and only weak transactivating sequences from Apicomplexan proteins have been reported (35). Because no trans-repressive domains have been reported to be functional in *Plasmodium*, we aimed to use CRISPRi with dSpCas9 alone to prevent or dampen association of RNAPII or other critical factors from binding to a specific gene.

For this approach, we constructed a CRISPRi plasmid similar to our 1^st^ generation CRISPR plasmid but without an HDR template, which is unnecessary for this application (Fig. 1). This plasmid encodes a nuclease dead dSpCas9::1xHA endonuclease (D10A, H840A), which permits the detection of GFP tags fused to a protein-of-interest to detect changes in gene expression. To test our CRISPRi plasmid, we utilized the marker-less PyALBA4::GFP transgenic parasite line, which was enriched by FACS to remove wild-type parasites and cloned by limiting dilution. Upon curing the CRISPR-RGR plasmid, these parasites regained pyrimethamine sensitivity, thus permitting introduction of another plasmid. We chose this parasite line as 1) the fusion of GFP to the C-terminus of PyALBA4 has no effect upon parasite growth and transmission, 2) is dispensable to asexual blood stage parasites, and thus 3) could allow direct testing of CRISPRi upon an endogenous gene and its gene control region. This final point is pertinent, as exogenous reporter expression cassette that have not performed well in predicting gene regulation effects upon endogenous genes in other systems.

Using RNA-seq data to estimate the 5’ UTR of the PyALBA4 locus (Fig. 4A, Supp. Fig. 3), we selected eleven individual sgRNA targets between the start of the contiguous RNA-seq reads (approximately −800) and the ATG. We selected sgRNAs that will target either the template or non-template strands of DNA to assess if strand-specific effects occur in *Plasmodium*, and that target DNA at various distances away from the ATG to assess positional effects (Fig. 4A). The nomenclature used for sgRNAs is based upon the distance of the N of the -NGG PAM sequence (e.g. −770) compared to the A of the ATG translational start codon (+1). As an experimental control, we also included a “no-target” sgRNA control, consisting of conserved sequence found in mCherry, mOrange and Tomato fluorescence markers (COT control) that are absent in this parasite. These twelve sgRNAs were cloned into independent CRISPRi plasmids and each was transfected into the marker-less PyALBA4::GFP parasite line.

**Figure 4:**
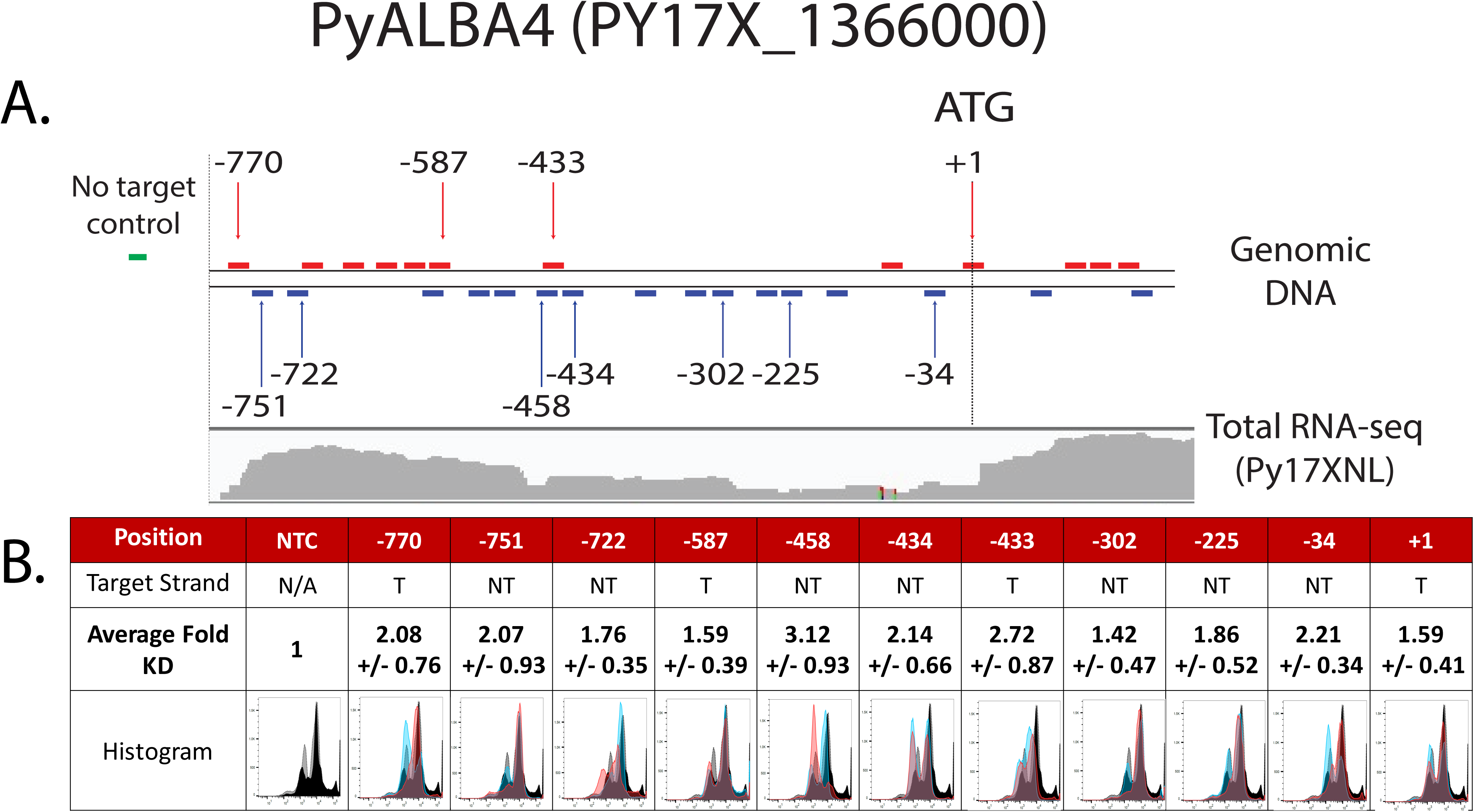
CRISPR interference of PyALBA4::GFP expression is dependent on the sgRNA target location. A.) Top: A schematic representation of the region upstream of the *pyalba4* coding sequence with all possible sgRNAs indicated by colored bars (red on template strand, blue on non-template strand). The 11 sgRNAs used for CRISPRi are indicated by arrows, with their numerical names denoting the position of the -N of the -NGG PAM sequence in reference to the ATG. Bottom: RNA-seq reads of Py17XNL parasites show the contiguous RNA-seq reads upstream of the *pyalba4* coding sequence. B.) Each sgRNA was tested in biological triplicate, with 2 technical replicates each and compared to the no-target control sgRNA. The average-fold knock down and standard error of the mean were calculated and combined across all replicates. Histograms are representative images (all from biological replicate two), which represent the GFP intensities for two technical replicates of the no-target control sgRNA (black and gray) and the on-target experimental sgRNA (red and blue).

To assess whether we could impose CRISPRi regulation upon this locus, we used flow cytometry to monitor PyALBA4::GFP protein abundance. This was measured in dSpCas9::1xHA-positive parasites expressing one of the eleven on-target sgRNAs or the no target sgRNA control that have the capacity for CRISPRi (Fig. 4, Supp Fig. 2). Parasites were synchronized to the schizont stage to minimize the effect of stage-specific variance in PyALBA4::GFP expression upon these observations. Parasites with background levels (negative or low) of dSpCas9::1xHA did not have a significant change in the median fluorescence intensity (MFI) of PyALBA4::GFP when using any of the on-target sgRNAs when compared to the no target sgRNA control (Supp. Fig. 2). In contrast, parasites expressing dSpCas9::1xHA above background produced a 2-3 fold reduction in PyALBA4::GFP signal when one of 6 of the 11 on-target sgRNAs was expressed (Fig. 4B, maximum knockdown was 3.12 fold). The two sgRNAs that produced the largest knockdown effect overlap the same sequence by 17nt, with one (−458) targeting the non-template strand and the other (−433) targeting the template strand. This indicates that transcription of *pyalba4* is particularly sensitive to the binding of dSpCas9 at this position. Comparison of this region’s sequence to known transcriptional effectors of the *P. falciparum* ApiAP2 family, we found that the core motif of AP2-G (PF3D7_1222600), GTAC, is present in the seed sequence and PAM site of these two sgRNAs although the functional significance of this remains to be seen (36). We did not observe any significant strand-dependent effects (average of 1.99-fold knock down for 4 template strand targets, average of 2.08-fold knock down for non-template strand targets). Finally, due to the presence of many auto-fluorescent events that could not be robustly gated out, we anticipate that we are underreporting the extent of knock down achieved. Taken together, these data show that CRISPRi with individual sgRNAs can modestly reduce expression of an endogenous gene in *Plasmodium yoelii*. These data further underscore the strong need to identify and characterize transcriptional activation and repression domains in *Plasmodium* to improve the scale of regulation possible by CRISPRi. This system provides an excellent platform on which to build those studies.

## Discussion

Here we present CRISPR-RGR, a ribozyme-based CRISPR system for *Plasmodium yoelii* that allows rapid and efficient gene editing in rodent-infectious parasites. This approach is powerful and provides advantages over current methods. First, the production of multiple sgRNAs from a single transcript greatly reduces the potential size of plasmids required for CRISPR-based gene editing by using a single promoter and terminator for sgRNA expression. Importantly, these sgRNAs can be designed to target a single gene in multiple locations as used here, but can be extended to target multiple genes. The utility of using multiple sgRNAs for a single gene is evident from the data presented here: the use of one sgRNA can result in a mixture of gene editing outcomes (plasmid integration and locus replacement), whereas the use of two sgRNAs produced only the desired locus replacement result. We strongly suggest that two or more sgRNAs be used for CRISPR-based gene editing where locus replacement is the desired outcome to eliminate the need for negative selection. Second, CRISPR-RGR can be programmed using synthetically produced DNA fragments that can encode the RGR, HDR, or both elements, which can be inserted in one or two molecular cloning steps. With the anticipated decreases in cost and increases in capacity to synthesize large DNA fragments, this approach will streamline reagent preparation and enable CRISPR screens at scale. Lastly, the FACS-based selection of transgenic parasites expressing SpCas9::GFP or a protein-of-interest fused to GFP reduces the number of mice required to produce transgenic parasite lines free from observable wild-type parasites. Because it is still possible that wild-type parasites remain in the population at levels lower than the limit of detection of PCR, caution is urged when phenotyping these parasites and it is our opinion that these parasites should be cloned prior to these studies.

CRISPR-RGR has a range of gene editing efficacies (50-100% editing) and time requirements that are comparable to other CRISPR systems in *Plasmodium* (11, 12, 18). However, we could rapidly isolate transgenic parasite populations (as per genotyping PCR) by selecting *P. yoelii* parasites that are expressing SpCas9::GFP by FACS. Coupled with the use of two sgRNAs to produce only locus replacement gene editing events, thus obviating the need for negative selection, CRISPR-RGR can generate the desired transgenic parasites quickly and with a minimum number of mice. Importantly, for subsequent genome modifications to be done, it is essential for these parasites to regain drug sensitivity so that another plasmid can be introduced. CRISPR-RGR achieves this effectively upon curing of the original plasmid.

Additionally, we demonstrate that CRISPR interference is possible in *Plasmodium yoelii* simply by binding dSpCas9 to the upstream gene control region of an endogenous gene, *pyalba4*. In contrast to early reports of the “rules” of CRISPRi in prokaryotes, model eukaryotes, and human cells, we did not observe a strand-specific effect on the knock down efficiency of dSpCas9. However, we did observe a positional effect, with the maximum effect of dSpCas9 binding approximately 430 to 450bp upstream of the translational start site. This positional effect was further corroborated by observing a similar knockdown effect when using two sgRNAs with overlapping target sequences (17 of 20 nucleotides). A sgRNA (−434) that causes dSpCas9 to be targeted adjacent to the site targeted by −458 and −434 produced a 2.14-fold knock down as well. While a 3.12-fold maximum knock down was observed, it would be advantageous to achieve much higher regulatory control. This will presumably require the identification of transcriptional activation and transcriptional repression domains that can be fused to dSpCas9 and placed in optimal genomic positions for activation (*e.g.* where little/no knockdown is observed when dSpCas9 alone binds) or repression (*e.g.* where maximum knock down is observed with dSpCas9 alone bound).

Based upon these findings, there are several improvements that can be made to improve CRISPR-based gene editing and regulation, some of which can be addressed using CRISPR-RGR. First, the transfection efficiency of all *Plasmodium* species is woefully low (e.g. 0.01 to 0.05%) compared to other eukaryotes (often >70%), even with the use of Amaxa nucleofectors (37). However, it is notable that these efficiencies are typically determined by the number of parasites receiving a plasmid and expressing some marker from it (e.g. GFP), and thus reflects the practical transfection efficiency for most applications. Work from the DeRisi lab demonstrated that the transfection of proteins into *P. falciparum* is much more efficient, with up to 1.7% of parasites receiving a Cas9::RFP protein (38). Thus, if delivery of SpCas9 bound to sgRNAs along with a HDR template can be achieved, plasmid-based gene editing may become unnecessary. Second, many genes-of-interest are essential to asexual blood stage parasites and thus cannot be deleted in this stage using conventional approaches. Instead, recombination-based approaches with stage-specific promoters can enable their excision at later time points, as long as these promoter are not sufficiently leaky or activate prematurely. While current CRISPR methodologies could use stage-specific promoters to drive SpCas9 expression, the leakiness of a single promoter could present similar problems. Using CRISPR-RGR, two discrete RNAPII promoters can be utilized to express the RGR and SpCas9 components, thus reducing effects of leaky expression by one of the promoters. Third, the range of efficiencies of sgRNAs is large and hard to predict computationally. Recent work by the Desai laboratory has completed a meta-analysis of a large number of sgRNAs that they have used in *P. falciparum*, and identified parameters that correlate with higher specificities and on-target efficiencies (39). An alternative approach would be to use multiple sgRNAs for each gene editing attempt, which can be achieved by CRISPR-RGR and appropriately designed HDR templates. Lastly, a similar strategy of using multiple sgRNAs can be applied to CRISPRi to avoid the preliminary assessment of the best placement of dSpCas9 in the gene control region. Coupled with effective trans-activation or trans-repression domains, or epigenetic regulators, we hypothesize that substantial regulatory effects can be achieved using CRISPR-RGR.

## Acknowledgements

We would like to thank Manuel Llinás and the members of the Lindner and Llinás labs, as well as Istvan Albert, Brian Dawson, and Howard Salis for critical discussions and technical assistance for this work. We also would like to thank our animal caretakers, and the Penn State Genomics Core Facility and Flow Cytometry Facility – University Park, PA. Funding for this work was supported by NIH grants R01AI123341 and R21AI130692 as well as by Pennsylvania State University start-up funds to SEL, and a Huck Institutes of the Life Sciences Graduate Research Innovation award to MPW. The funders had no role in the study design, data collection and interpretation, or the decision to submit the work for publication.

## Materials and Methods

### Ethics

All animal handling followed the Association and Accreditation of Laboratory Animal Care (AAALAC) guidelines and these protocols were approved by the Pennsylvania State University Institutional Animal Care and Use Committee, #42678-01. In addition, all procedures with vertebrate animals were conducted in accordance with the Guide for Care and Use of Laboratory Animals of the National Institutes of Health with approved Office for Laboratory Animal Welfare (OLAW) assurance.

### Experimental Animals and Parasite Lines

Six to eight-week-old female Swiss Webster (SW) mice were obtained from Envigo and used for all experiments described. *Plasmodium yoelii* 17XNL non-lethal parasites and *Plasmodium falciparum* 3D7 strain parasites were used for all experiments.

### Plasmid Construction

A step-by-step guide for creating RGR-based sgRNAs for *Plasmodium* is provided as Supp File 1. The CRISPR editing and CRISPR interference plasmids (illustrated in Fig. 1 and Supp Fig. 1, with sequences provided in Supp File 2.) were constructed from a combination of conventional molecular cloning and gene synthesis (GeneWiz) approaches. *S. pyogenes* Cas9::GFP and dCas9 coding sequences were obtained from Addgene (Plasmids #48138, #42335) and were sub-cloned into plasmid pSL0444 containing the HsDHFR drug cassette (primer sequences listed in Supp Table 1). An empty cassette consisting of the *pbef1α* promoter and *pbdhfr*3’UTR were synthesized (GeneWiz) and cloned into the HsDHFR+Cas9::GFP and HsDHFR+dCas9 plasmids and used for expression of synthetic hammerhead Ribozyme-Guide-Hepatitis Delta Virus Ribozyme (RGR) transcripts. RGR coding sequences were synthesized (GeneWiz, sgRNA target sequences provided in Supp Table 3) based on the “optimized” sgRNA structure described in (30). Homology-directed repair (HDR) templates required for gene deletions and gene insertions (mediated by the editing plasmids) were generated by SOE PCR amplification of *P. yoelii* genomic DNA with oligos specific to the *pyalba4* locus. In addition, exogenous barcode sequences (gene deletions) or a GFPmut2 tag (gene insertion) were included between the two arms of the HDR template. The second-generation editing plasmids were constructed by synthesizing two portions of the first-generation plasmid with different promoters and 3’UTRs to reduce the possibility of recombination events. A plasmid (pSL1156) containing the *pyef1α* 3’UTR, a multiple cloning site, and the *pyalba4* 191bp HDR template was used to move these sequences into the first-generation CRISPR editing plasmid (pSL0999) via XbaI and NheI. A DNA fragment including *pybip* 3’UTR and *pydhfr* promoter regions were then cloned into the AgeI and SpeI sites found between the RGR and Cas9 sequences. The penultimate construction (pSL1165) has no repetitive or redundant elements controlling SpCas9, RGR, and HsDHFR expression cassettes, with a multiple cloning site for insertion of unique promoters to drive RGR expression. Candidate promoters for the RGR cassette were generated by PCR amplifying 1.5kb of the sequences upstream from (and including) the ATG of *pybip* (PY17X_0822200) and *pygapdh* (PY17X_1330200) with NheI and NotI RE sites on the 5’ and 3’ ends respectively (primer sequences listed in Supp Table 2). These products were inserted into and sequenced in pCR-Blunt, and then subcloned into pSL1165 to create pSL1166 (using the *pybip* promoter) and pSL1211 (using the *pygapdh* promoter). These plasmids were transfected into *Plasmodium yoelii* 17XNL strain parasites and were analyzed for their editing efficiency. The empty vector and *pyalba4*-targeted plasmids described in this work will be available on Addgene.

### Generation of Transgenic *P. yoelii* and *P. falciparum* Parasites

Transgenic *P. yoelii* parasites were generated as previously described (2, 40) but with constant pyrimethamine selection post-transfection. Presence of transgenic parasites was confirmed by genotyping PCR using primer with sequences external to the targeting sequences (Supp Table 1).

### Fluorescence-Activated Cell Sorting (FACS)

Synchronized and Accudenz-purified *P. yoelii* schizonts (2, 40) were sorted on a Beckman Coulter MoFlo Astrios EQ Cell Sorter for SpCas9::GFP or PyALBA4::GFP expression above background (determined using uninfected blood as a negative control). For each replicate, 5,000 to 10,000 GFP-positive events were sorted and IV injected into a naïve mouse. Pyrimethamine pressure was retained until parasitemia reached ∼1% in the mouse. Parasite populations were assessed by genotyping PCR.

### Curing the CRISPR:RGR Plasmid from PyALBA4::GFP Parasites

Sorted PyALBA4::GFP parasites were injected intravenously into a naïve mouse and parasites developed for 9-10 days. Once ∼1% parasitemia was reached, these parasites were cloned out by limited dilution to achieve a clonal population of PyALBA4::GFP parasites. Naïve mice were infected with these parasites for 4 days, and then were treated with pyrimethamine-drugged water to test drug sensitivity.

### Exflagellation Measurements

Male gametocyte activation was determined by counting the number of exflagellation events in ten 40x magnification phase contrast fields (Leica ICC50 HD) in a monolayer of blood cells obtained from a tail snip. These measurements were taken daily for 18 days post infection with either wild-type or transgenic parasite lines. Three biological replicates were performed for each line, each with one mouse.

### Live Fluorescence and ImmunoFluorescence Assays (IFAs)

PyALBA4::GFP-expressing parasites were imaged by both live fluorescence and indirect immunofluorescence assays. Live fluorescence and IFA micrographs were collected on a Zeiss fluorescence/phase contrast microscope (Zeiss Axioscope A1 with 8-bit AxioCam ICc1 camera) on a 63x objective. IFAs were performed as previously described (41) at room temperature with a two hour blocking step and one hour incubations with primary antibodies (mouse anti-GFP (DSHB-GFP-4C9-ds) and rabbit anti-ACP (Pocono Rabbit Farm and Laboratory, Custom), both diluted 1:1000) and secondary antibodies (anti-mouse and anti-rabbit, Alexa Fluor conjugated AF488 and AF594, respectively (Invitrogen, #A-11001 and #A-11012), both diluted 1:500). Images from both live fluorescence and IFAs were collected and processed with Zen imaging software (Zeiss).

### Flow Cytometry Assessment of CRISPR Interference

Schizonts were synchronized by overnight growth *in vitro* and purified by a discontinuous Accudenz density gradient (2, 40), and were prepared identically to IFA samples as described above. Samples were stained with rabbit anti-GFPmut2 (Pocono Rabbit Farm and Laboratory, #32180) bound by anti-rabbit IgG conjugated to AF488 (Invitrogen, #A-11008) and mouse anti-HA (Clone 12CA5) bound by an anti-mouse IgG conjugated to AF405 (Invitrogen, #A-31553). Samples were analyzed on a BD-LSR Fortessa using uninfected RBCs, RBCs infected with Py17XNL wild-type parasites or PyALBA4::GFP-expressing parasites as negative and positive controls. Gating was established using samples stained with a single primary/secondary for each channel. Data was processed using FlowJo (v10.4.1) to determine the change in median fluorescence intensity (MFI) of GFP expression in parasites expressing dSpCas9::1xHA. MFIs were averaged within each biological replicate and fold-change for each experimental sgRNA compared to the no-target control was calculated. The fold-change values were averaged across biological replicates for each sgRNA target.

**Supplemental Figure 1: The 2^nd^ Generation editing plasmids with different constitutive promoters for the transcription of the RGR element show efficient editing of the *pyalba4* locus.** The *pybip* (A) and *pygapdh* (B) promoters were PCR amplified and cloned into the unique NotI and NheI sites to drive expression of the RGR element used to delete the *pyalba4* coding sequence. Each plasmid was transfected into *Plasmodium yoelii* 17XNL parasites, and edited parasites were detected by genotyping PCR. Parasites transfected with the plasmid using the *pybip* promoter for RGR expression (A) were enriched via FACS for SpCas9::GFP expression. Sorted SpCas9::GFP parasites were genotyped by PCR, revealing that only transgenic parasites are observed.

**Supplemental Figure 2: Flow cytometric data of CRISPRi experiments.** Three biological replicates, each with two technical replicates, were performed with the 11 sgRNA targets and the no target control, each with negative controls. Gating of SpCas9::HA expressing parasites was determined by the HA-only antibody staining of the no target control replicates (set at 2×10^2 for each replicate). HA-positive and HA-negative were gated separately and their GFP intensities are displayed in each histogram. Values for each of these replicates can be found in Supplemental Table 1.

**Supplemental Figure 3: RNA-seq read coverage of the region upstream of the *pyalba4* coding sequence.** Three replicates of Py17XNL WT RNA-seq were performed and alignments to the *pyalba4* locus show reads aligning across ∼800bp of sequence upstream of the ATG. These mapped reads provide a basis for selecting sgRNAs for the CRISPRi experiments. Dashes (top) indicate 100bp of sequence.

**Supplemental Table 1: Oligonucleotide sequences used in this study.** These sequences include primers used to amplify the CRISPR plasmid elements, such as the SpCas9, promoter sequences, and the HDR templates used, as well as those used for genotyping PCR of the transgenic parasites that were generated.

**Supplemental Table 2: Complete CRISPRi Flow Cytometry Analysis** Individual CRISPRi flow cytometry values for each replicate performed in this study are provided, including all median fluorescence intensities for both HA-positive and HA-negative populations.

**Supplemental Table 3: Nucleotide sequences and positions of sgRNAs used in this study** Each 20nt sequence of the sgRNA targets used in this study are annotated with their position and the experiments they were used for.

**Supplemental File 1: A step-by-step tutorial for creation of sgRNAs by the Ribozyme-Guide-Ribozyme (RGR) methodology for *Plasmodium*.**

**Supplemental File 2: Descriptions and complete sequences of plasmids used in this study**

## References

1. Koning-Ward TFd, and CJJ, Waters AP. 2000. The Development of Genetic Tools for Dissecting the Biology of Malaria Parasites. Annual Review of Microbiology 54:157–185.

2. Jongco AM, Ting L-M, Thathy V, Mota MM, Kim K. 2006. Improved transfection and new selectable markers for the rodent malaria parasite Plasmodium yoelii. Molecular and Biochemical Parasitology 146:242–250.

3. Janse CJ, Ramesar J, Waters AP. 2006. High-efficiency transfection and drug selection of genetically transformed blood stages of the rodent malaria parasite Plasmodium berghei. Nature Protocols 1:346.

4. Reece SE, Thompson J. 2008. Transformation of the rodent malaria parasite Plasmodium chabaudi and generation of a stable fluorescent line PcGFPCON. Malaria journal 7:183–183.

5. Lin J-w, Annoura T, Sajid M, Chevalley-Maurel S, Ramesar J, Klop O, Franke-Fayard BMD, Janse CJ, Khan SM. 2011. A Novel ‘Gene Insertion/Marker Out’ (GIMO) Method for Transgene Expression and Gene Complementation in Rodent Malaria Parasites. PLOS ONE 6:e29289.

6. Lacroix C, Giovannini D, Combe A, Bargieri DY, Spath S, Panchal D, Tawk L, Thiberge S, Carvalho TG, Barale JC, Bhanot P, Menard R. 2011. FLP/FRT-mediated conditional mutagenesis in pre-erythrocytic stages of Plasmodium berghei. Nat Protoc 6:1412–28.

7. Collins CR, Das S, Wong EH, Andenmatten N, Stallmach R, Hackett F, Herman J-P, Müller S, Meissner M, Blackman MJ. 2013. Robust inducible Cre recombinase activity in the human malaria parasite Plasmodium falciparum enables efficient gene deletion within a single asexual erythrocytic growth cycle. Molecular Microbiology 88:687–701.

8. Schwach F, Bushell E, Gomes AR, Anar B, Girling G, Herd C, Rayner JC, Billker O. 2015. PlasmoGEM, a database supporting a community resource for large-scale experimental genetics in malaria parasites. Nucleic Acids Research 43:D1176–D1182.

9. Fonager J, Franke-Fayard BM, Adams JH, Ramesar J, Klop O, Khan SM, Janse CJ, Waters AP. 2011. Development of the piggyBac transposable system for Plasmodium berghei and its application for random mutagenesis in malaria parasites. BMC Genomics 12:155.

10. Muñoz EE, Hart KJ, Walker MP, Kennedy MF, Shipley MM, Lindner SE. 2017. ALBA4 modulates its stage-specific interactions and specific mRNA fates during Plasmodium yoelii growth and transmission. Molecular Microbiology 106:266–284.

11. Ghorbal M, Gorman M, Macpherson CR, Martins RM, Scherf A, Lopez-Rubio JJ. 2014. Genome editing in the human malaria parasite Plasmodium falciparum using the CRISPR-Cas9 system. Nat Biotechnol 32:819–21.

12. Wagner JC, Platt RJ, Goldfless SJ, Zhang F, Niles JC. 2014. Efficient CRISPR-Cas9-mediated genome editing in Plasmodium falciparum. Nat Methods 11:915–8.

13. Zhang C, Xiao B, Jiang Y, Zhao Y, Li Z, Gao H, Ling Y, Wei J, Li S, Lu M, Su XZ, Cui H, Yuan J. 2014. Efficient editing of malaria parasite genome using the CRISPR/Cas9 system. MBio 5:e01414–14.

14. Aly ASI, Vaughan AM, Kappe SHI. 2009. Malaria Parasite Development in the Mosquito and Infection of the Mammalian Host. Annual Review of Microbiology 63:195–221.

15. Butler Noah S, Schmidt Nathan W, Vaughan Ashley M, Aly Ahmed S, Kappe Stefan HI, Harty John T. 2011. Superior Antimalarial Immunity after Vaccination with Late Liver Stage-Arresting Genetically Attenuated Parasites. Cell Host & Microbe 9:451–462.

16. Kirkman LA, Lawrence EA, Deitsch KW. 2014. Malaria parasites utilize both homologous recombination and alternative end joining pathways to maintain genome integrity. Nucleic Acids Research 42:370–379.

17. Lee AH, Symington LS, Fidock DA. 2014. DNA Repair Mechanisms and Their Biological Roles in the Malaria Parasite Plasmodium falciparum. Microbiology and Molecular Biology Reviews 78:469–486.

18. Zhang C, Gao H, Yang Z, Jiang Y, Li Z, Wang X, Xiao B, Su XZ, Cui H, Yuan J. 2017. CRISPR/Cas9 mediated sequential editing of genes critical for ookinete motility in Plasmodium yoelii. Mol Biochem Parasitol 212:1–8.

19. Liu C, Li Z, Jiang Y, Cui H, Yuan J. 2018. Generation of Plasmodium yoelii malaria parasite carrying double fluorescence reporters in gametocytes. Molecular and Biochemical Parasitology 224:37– 43.

20. Qian P, Wang X, Yang Z, Li Z, Gao H, Su X-z, Cui H, Yuan J. 2018. A Cas9 transgenic Plasmodium yoelii parasite for efficient gene editing. Molecular and Biochemical Parasitology 222:21–28.

21. Zhang C, Li Z, Cui H, Jiang Y, Yang Z, Wang X, Gao H, Liu C, Zhang S, Su X-z, Yuan J. 2017. Systematic CRISPR-Cas9-Mediated Modifications of Plasmodium yoelii ApiAP2 Genes Reveal Functional Insights into Parasite Development. mBio 8.

22. Baum J, Papenfuss AT, Mair GR, Janse CJ, Vlachou D, Waters AP, Cowman AF, Crabb BS, de Koning-Ward TF. 2009. Molecular genetics and comparative genomics reveal RNAi is not functional in malaria parasites. Nucleic Acids Research 37:3788–3798.

23. Prommana P, Uthaipibull C, Wongsombat C, Kamchonwongpaisan S, Yuthavong Y, Knuepfer E, Holder AA, Shaw PJ. 2013. Inducible Knockdown of Plasmodium Gene Expression Using the glmS Ribozyme. PLOS ONE 8:e73783.

24. Russo I, Oksman A, Vaupel B, Goldberg DE. 2009. A calpain unique to alveolates is essential in Plasmodium falciparum and its knockdown reveals an involvement in pre-S-phase development. Proceedings of the National Academy of Sciences of the United States of America 106:1554– 1559.

25. Muralidharan V, Oksman A, Iwamoto M, Wandless TJ, Goldberg DE. 2011. Asparagine repeat function in a Plasmodium falciparum protein assessed via a regulatable fluorescent affinity tag. Proceedings of the National Academy of Sciences of the United States of America 108:4411– 4416.

26. Ganesan SM, Falla A, Goldfless SJ, Nasamu AS, Niles JC. 2016. Synthetic RNA-protein modules integrated with native translation mechanisms to control gene expression in malaria parasites. Nature communications. 7(10727. doi:10.1038/ncomms10727.

27. Gao Y, Zhao Y. 2014. Self-processing of ribozyme-flanked RNAs into guide RNAs in vitro and in vivo for CRISPR-mediated genome editing. J Integr Plant Biol 56:343–9.

28. Zhang W-W, Matlashewski G. 2015. CRISPR-Cas9-Mediated Genome Editing in Leishmania donovani. mBio 6.

29. Lee RTH, Ng ASM, Ingham PW. 2016. Ribozyme Mediated gRNA Generation for In Vitro and In Vivo CRISPR/Cas9 Mutagenesis. PLOS ONE 11:e0166020.

30. Dang Y, Jia G, Choi J, Ma H, Anaya E, Ye C, Shankar P, Wu H. 2015. Optimizing sgRNA structure to improve CRISPR-Cas9 knockout efficiency. Genome Biol 16:280.

31. Peng D, Tarleton R. 2015. EuPaGDT: a web tool tailored to design CRISPR guide RNAs for eukaryotic pathogens. Microbial Genomics 1:-.

32. Qi LS, Larson MH, Gilbert LA, Doudna JA, Weissman JS, Arkin AP, Lim WA. 2013. Repurposing CRISPR as an RNA-guided platform for sequence-specific control of gene expression. Cell 152:1173–83.

33. Larson MH, Gilbert LA, Wang X, Lim WA, Weissman JS, Qi LS. 2013. CRISPR interference (CRISPRi) for sequence-specific control of gene expression. Nature Protocols 8:2180.

34. Maeder ML, Linder SJ, Cascio VM, Fu Y, Ho QH, Joung JK. 2013. CRISPR RNA–guided activation of endogenous human genes. Nature Methods 10:977.

35. Pino P, Sebastian S, Kim EA, Bush E, Brochet M, Volkmann K, Kozlowski E, Llinas M, Billker O, Soldati-Favre D. 2012. A tetracycline-repressible transactivator system to study essential genes in malaria parasites. Cell Host Microbe 12:824–34.

36. Campbell TL, De Silva EK, Olszewski KL, Elemento O, Llinás M. 2010. Identification and Genome-Wide Prediction of DNA Binding Specificities for the ApiAP2 Family of Regulators from the Malaria Parasite. PLOS Pathogens 6:e1001165.

37. Janse CJ, Franke-Fayard B, Mair GR, Ramesar J, Thiel C, Engelmann S, Matuschewski K, Gemert GJv, Sauerwein RW, Waters AP. 2006. High efficiency transfection of Plasmodium berghei facilitates novel selection procedures. Molecular and Biochemical Parasitology 145:60–70.

38. Crawford ED, Quan J, Horst JA, Ebert D, Wu W, DeRisi JL. 2017. Plasmid-free CRISPR/Cas9 genome editing in Plasmodium falciparum confirms mutations conferring resistance to the dihydroisoquinolone clinical candidate SJ733. PLOS ONE 12:e0178163.

39. Ribeiro JM, Garriga M, Potchen N, Crater AK, Gupta A, Ito D, Desai SA. 2018. Guide RNA selection for CRISPR-Cas9 transfections in Plasmodium falciparum. International Journal for Parasitology 48:825–832.

40. Lindner SE, Mikolajczak SA, Vaughan AM, Moon W, Joyce BR, Sullivan WJ, Jr., Kappe SH. 2013. Perturbations of Plasmodium Puf2 expression and RNA-seq of Puf2-deficient sporozoites reveal a critical role in maintaining RNA homeostasis and parasite transmissibility. Cell Microbiol 15:1266–83.

41. Miller JL, Harupa A, Kappe SHI, Mikolajczak SA. 2012. Plasmodium yoelii Macrophage Migration Inhibitory Factor Is Necessary for Efficient Liver-Stage Development. Infection and Immunity 80:1399–1407.

